# Signature Distance: Generalizing Energy Statistics

**DOI:** 10.1101/2024.10.23.619602

**Authors:** Nicolo’ Lazzaro, Raffaele Marchesi, Gianluca Leonardi, Jacopo Tessadori, Marco Chierici, Monica Moroni, Gabriele Sales, Toma Tebaldi, Giuseppe Jurman

## Abstract

Comparing empirical distributions is central to generative model evaluation, hypothesis testing and data augmentation in high-dimensional biological data. Established methods such as energy distance summarize each point’s relationship to the opposing distribution through a single expected distance, providing sensitivity to location shifts. We introduce Signature Distance (SD), a statistical distance that compares empirical distributions through the mean absolute difference of their sorted pointwise distance profiles. SD is a structural generalization of energy distance and matches its quadratic pairwise-distance cost, with an additional sorting step. In controlled experiments and on TCGA pan-cancer transcriptomic data, we show that (1) SD detects density changes with greater sensitivity than energy distance in the tested scale-contraction scenarios; (2) per-point mean-distance and signature-profile landscapes reveal the geometric mechanisms behind their different penalties; (3) linearly interpolated biological samples that receive no increased penalty from energy distance are penalized by SD; (4) SD provides a direct differentiable potential energy for model-free Langevin data expansion, with a bootstrap resampling protocol to assess the stopping epoch; and (5) SD is directly usable as a differentiable generative training loss. Code to reproduce all experiments is available at https://github.com/lazzaronico/signature-distance.

## Introduction

Comparing empirical distributions is a recurring problem in computational biology: assessing whether synthetic data resembles real data (Luecken and Theis, 2019), testing for differences between treatment and control groups, and guiding iterative data generation. In high-dimensional settings, where biological measurements routinely span hundreds to thousands of features, distances between points concentrate (Beyer et al., 1999), and standard comparison methods lose discriminative power.

A widely adopted family of methods relies on expected pairwise distances. Energy distance (ED; Székely and Rizzo (2013)) compares distributions through the difference between the expected cross-distance and the expected within-set distances, yielding a raw statistic whose square root embeds in a Hilbert space (a complete inner-product space) and supports permutation-based hypothesis testing. Related methods share the same scalar-reduction structure: maximum mean discrepancy (MMD; Gretton et al. (2012)) generalises ED to arbitrary reproducing kernels (distance-induced kernels recover energy statistics; Sejdinovic et al. 2013), while the Fréchet Inception Distance (FID; Heusel et al. (2017)) compares Gaussian-fitted moments in a learned feature space. We use ED as the representative baseline throughout.

Because this scalar reduction captures only the mean pairwise distance, it is well-suited to detecting global location shifts, while reducing each point’s radial profile to its mean. Two distributions can have identical expected cross-distances while differing in density, shape, or internal manifold organization. As we show in Section 3.1, controlled density perturbations are detected more reliably by SD than by ED at a limited sample size.

At the other end of the computational spectrum, the full-dimensional 1-Wasserstein distance (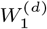; Villani 2009; Peyré and Cuturi 2019) provides a principled framework for geometric distributional comparison, but the cubic cost of exact equal-mass assignment can be prohibitive at the sample sizes common in omics data. The superscript denotes the ambient dimension; the subscript denotes the transport order. Topological data analysis (Carlsson, 2009; Edelsbrunner and Harer, 2010) captures multi-scale geometric structure via filtrations but produces summaries that are expensive to compare natively.

We introduce Signature Distance, which bridges this gap. A point’s sorted distances to all members of a population form a one-dimensional fingerprint of its local neighborhood: the nearest neighbors determine the front of the sorted array, while the distant points fill its tail. Because this fingerprint encodes the full radial density profile, it captures structural features such as density gradients, cluster boundaries and manifold curvature that are lost when neighborhood relationships are collapsed to a scalar mean. SD compares these fingerprints across two populations by computing the *W*_1_ distance between each point’s intra- and cross-distribution sorted profiles, then sums the average discrepancies over the two reference populations (Figure 1; see Methods, Section 2.1). This decomposition into one-dimensional *W*_1_ problems matches ED’s quadratic pairwise-distance cost, with an additional sorting step, while retaining sensitivity to local density structure.

**Fig. 1:**
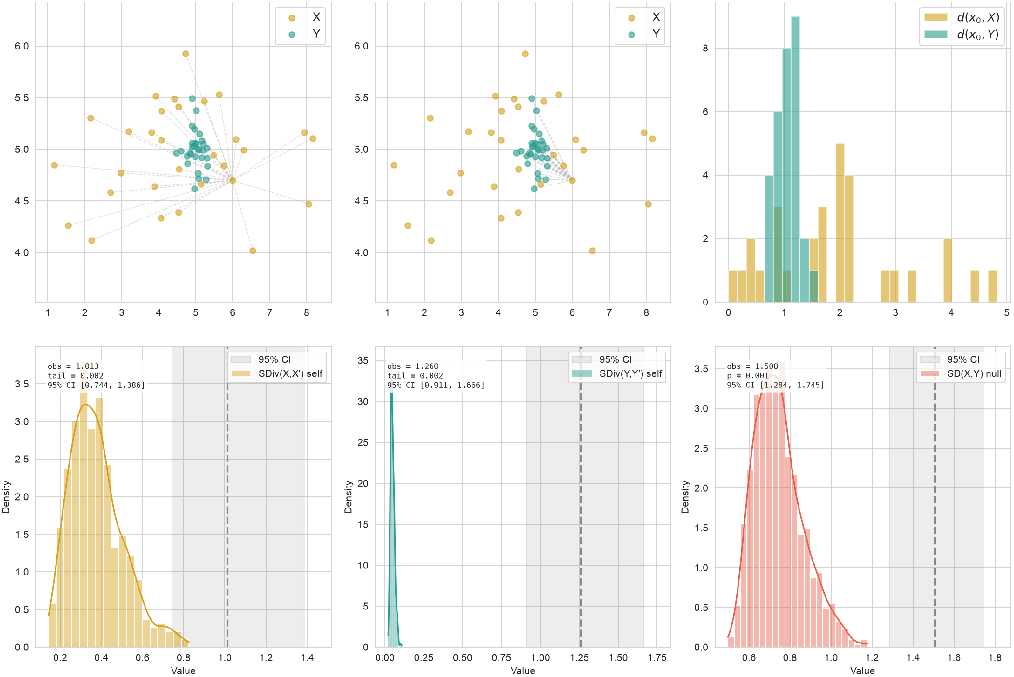
The geometric signature and its use as a two-sample test. Two populations *X* (gold, dispersed) and *Y* (teal, compact cluster) are drawn from different distributions in ℝ^2^. *Top left:* the signature of a reference point *x*_0_ ∈ *X* consists of its sorted distances to all points in *X* (intra-distribution distances, dashed lines). *Top centre:* the same reference point’s distances to all points in *Y* (cross-distribution distances). *Top right:* histogram comparison: if *X* and *Y* were drawn from the same distribution, these two distance profiles would be statistically similar; their divergence (measured by sorted-rank L1 distance) quantifies how much the neighbourhood geometry around *x*_0_ differs between populations. *Bottom row:* self-bootstrap reference distributions (shaded) and observed statistics (dashed line) for both directional divergences SDiv(*X* ∥ *Y* ) and SDiv(*Y* ∥ *X*), and a permutation-test null for the symmetric SD(*X, Y* ). Both directional divergences exceed their self-bootstrap references; symmetric SD exceeds its permutation null (*p* = 0.001). The grey bands are 95% percentile bootstrap confidence intervals. Resampling settings are given in Section S2.

The present work makes five contributions. First, we formally define Signature Distance and establish its bounds and separation properties, analytically deriving its structural relationship to ED (Section 2.1). Second, we show that SD detects distributional differences with greater sensitivity than ED in the tested density-change settings, using controlled perturbations (Section 3.1). Third, we show that per-point mean-distance and signature-profile penalties reveal the geometric mechanisms behind their different behaviour (Section 3.2). Fourth, we show that linearly interpolated biological samples unpenalized by ED receive an increased penalty from SD, and that SD serves as a differentiable potential energy for model-free Langevin data expansion (Sections 3.3–3.4). Fifth, we show that SD is directly usable as a differentiable generative training loss, and introduce a glocal training protocol for multi-population settings that enables fair comparison of distributional losses on tissue-conditioned generation of TCGA landmark gene expression profiles (Section 3.5).

## Methods

### Signature Distance

Given two empirical point clouds *X* = {*x*_1_, …, *x*_*n*_} and *Y* = {*y*_1_, …, *y*_*m*_} in ℝ^*d*^, write *q*_*X*_ (*z, u*) for the *u*-quantile of the distances from a reference point *z* to the points in *X*, and similarly for *Y* . Here *z* is one of the observations supplying the reference point; 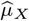 and 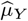 denote the uniform empirical distributions on *X* and *Y* . These quantiles describe the empirical radial-distance distributions; their discrete computation is given in Appendix C.

The signatures are compared by integrating the absolute difference of their quantiles. This operation is the one-dimensional 1-Wasserstein distance between the radial distributions:

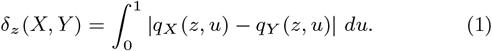

We define the raw signature statistic *S* as the sum of the two average pointwise divergences:

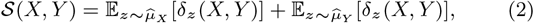

with 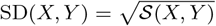.

In the population limit, when *X* and *Y* are drawn from the same distribution, each point’s intra- and cross-distribution distance profiles converge to the same CDF, yielding SD → 0; for finite samples, SD(*X, Y* ) ≈ 0 under the null, with the permutation test quantifying the typical null variation. When the distributions differ in density, shape, or location, the sorted distance profiles diverge.

#### Theoretical Properties

##### Identity of indiscernibles

All atoms, including zero self-distances, retain their empirical mass. When evaluating an identical set SD(*X, X*), the intra- and cross-signatures for every point are identical, so SD(*X, X*) = 0.

##### Properties

SD is nonnegative, symmetric and zero exactly for equal Euclidean probability measures with finite first moments (Appendix A).

##### Upper and lower bounds (Appendix A, Theorem 1)

The summed statistic *S* = SD^2^ is bounded below by the raw energy statistic ε and above by twice the exact 1-Wasserstein distance:

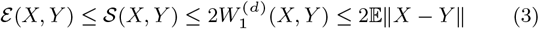

ε = 2 E ∥*X* − *Y* ∥ − E ∥*X* − *X*^′^∥ − E ∥*Y* − *Y* ^′^∥ is the raw energy expression; ED denotes its rooted metric 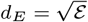 in the figures.

##### Scale-invariant normalization (SD_norm_)

The rightmost bound in the chain, 2E ∥*X* − *Y* ∥, provides a scale factor that normalises *S* into a unit interval. This permits the definition of an absolute, scale-invariant index:

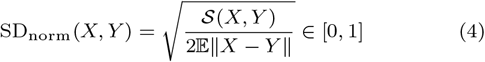

##### Computational complexity

Pairwise distances scale as *O*((*n* + *m*)^2^), and the per-point sorting scales as *O*((*n* + *m*) · max(*n, m*) log max(*n, m*)). For comparable sizes this gives *O*(*n*^2^*d*+ *n*^2^ log *n*) time; ED avoids the sorting term.

#### Asymmetric decomposition and computational efficiency

Equation 2 decomposes naturally into two directed components. In a generative setting where *X* denotes generated samples and *Y* real reference data, these components admit a precision/recall interpretation (Kynkäänniemi et al., 2019):

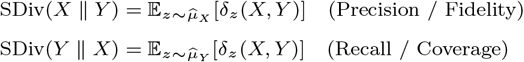

so that S(*X, Y* ) = SDiv(*X* ∥ *Y* ) + SDiv(*Y* ∥ *X*). SDiv(*X* ∥ *Y* ) (precision) evaluates whether each generated sample’s neighborhood structure matches that of the real manifold; SDiv(*Y* ∥ *X*) (recall) evaluates whether each real sample’s local neighborhood is faithfully reproduced by the generated distribution. This decomposition has a practical consequence for iterative algorithms: when *Y* is a fixed reference, the intra-reference radial quantiles *q*_*Y*_ (*y, u*) can be precomputed once and cached, avoiding repeated real–real distance and sorting work for the real-anchor component. These components differ from the independent precision and recall estimates reported in the Supporting Information.

#### Finite-sample inference and hypothesis testing

SD supports a standard permutation-based two-sample test. Given samples *X* (size *n*) and *Y* (size *m*), compute the observed statistic *d*_obs_ = SD(*X, Y* ). Pool both sets into *Z* = *X* ∪ *Y* and, for *b* = 1, …, *B* permutations, randomly partition *Z* into groups of size *n* and *m* and compute *d*_*b*_ = SD(*Z*_1:*n*_, *Z*_*n*+1:*n*+*m*_). The empirical p-value is 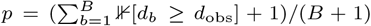, following the pseudocount convention of Phipson and Smyth (2010) to ensure strictly positive estimates and controlled Type I error. The test reuses the precomputed full pairwise distance matrix, permuting indices rather than recomputing distances, although recomputing signatures still requires sorting within each partition.

This permutation test evaluates Fisher’s conditional null: whether the observed partition produces a distance larger than expected under random label assignment, conditional on the specific sample at hand (Ernst, 2004). For exact enumeration, its resolution is bounded by the combinatorial space 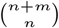, which for moderate samples (*n* = *m* = 15,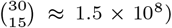) can yield valid p-values below 0.001. For a Monte Carlo test using *B* random permutations, the plus-one estimate instead has minimum 1*/*(*B* + 1). To complement the randomisation test with a measure of population-level stability, we provide bootstrap confidence intervals (Efron and Tibshirani, 1993): resampling the observed groups with replacement over *B*_boot_ iterations and recomputing SD yields an empirical estimate of the distance’s sampling distribution, whose 95% percentile interval summarizes variation across hypothetical replicate cohorts. Figure-specific settings are given in Section S2.

### Column Distance and density-level constraints

The signature matrix can be integrated along either axis, yielding complementary statistics. SD integrates *row-wise*, comparing per-point neighbourhood topology (Eq. 2). A natural complement integrates *column-wise*: for each quantile level *u*, the population distribution of radial quantiles should match across the two sets. We define the asymmetric Column Divergence and the **Squared Column Distance (CD^2^)**

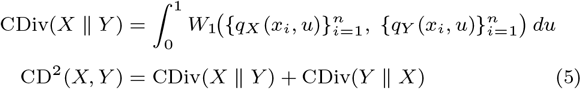

Because each column is one-dimensional, *W*_1_ reduces to sorting, with 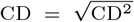. CD constrains the *global density level-set structure* (the fraction of points within each nearest-neighbour shell) while SD constrains the local topology. Combining the row and column signals via their Pythagorean (L2) combination gives:

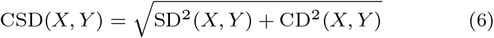

Column sorting marginalises spatial identity: CD does not track *which* points contribute to each density level.

### Grounded Signature Distance

SD compares each point’s radial neighbourhood profiles between the two distributions but does not enforce spatial correspondence with any specific reference point. CD addresses this through population-level density matching but marginalises point identity. Grounded Signature Distance (GSD) provides a more direct solution by grounding each point to its nearest reference neighbour. We define the asymmetric Grounding Divergence and the symmetric Squared Grounded Signature Distance:

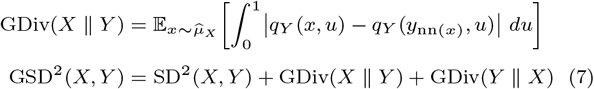

where *y*_nn(*x*)_ is the nearest neighbour in *Y* to *x*, and 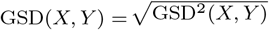. The grounding term evaluates whether each point’s distance profile matches that of its nearest neighbour in the opposing set, spatially grounding the profile comparison. This captures density-level information that CD addresses through column sorting, with the NN lookup and profile comparison reusing the already-computed distance and signature matrices. When used as a training loss, the NN targets are treated as fixed reference values (detached from the computational graph), providing a fixed spatial anchor for the gradient update.

## Results

Controlled experiments use synthetic two-dimensional distributions; all biological experiments use The Cancer Genome Atlas (TCGA) pan-cancer gene expression dataset (The Cancer Genome Atlas Research Network, 2008).

### Sensitivity to distributional differences

To illustrate the behaviour of SD, CSD, and ED under controlled distributional perturbations, we consider a two-dimensional Gaussian scenario (Figure 2). A reference distribution *X* of 50 points from a standard two-dimensional Gaussian is tested against three alternatives: same distribution, location shift, and density change. Each alternative is sampled independently.

**Fig. 2:**
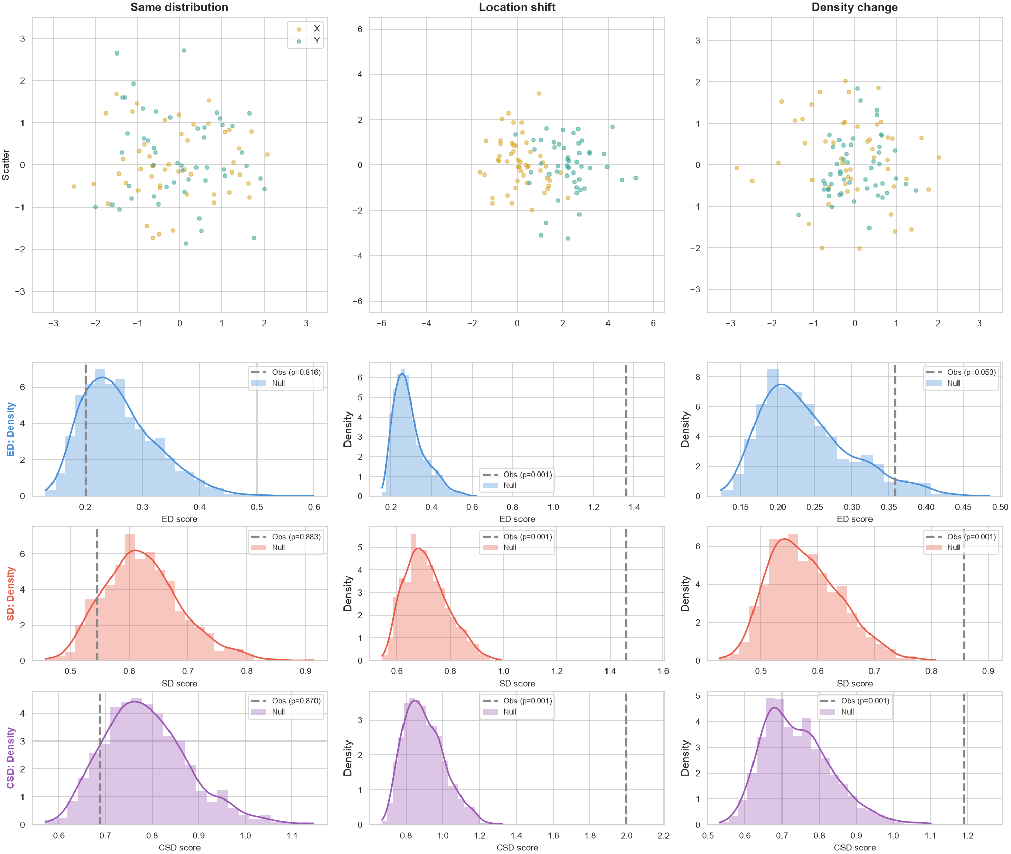
Sensitivity comparison of SD, CSD, and ED on controlled two-dimensional scenarios. *Top row:* scatter plots of *X* (gold) and *Y* (teal). *Rows 2–4:* permutation-test null distributions (shaded) and observed values (dashed lines) for ED, SD, and CSD. *Left (same distribution): X* and *Y* are drawn from the same Gaussian; all statistics correctly fall within their null. *Centre (location shift): Y* is shifted; all three detect the difference (*p* = 0.001). *Right (density change): Y* is drawn independently from a Gaussian with scale 0.7. SD and CSD detect the contraction (*p* = 0.001 each), while ED does not reject at the 0.05 level (*p* = 0.053). The first replicate of the *n* = 50 comparison is shown. Repeated comparisons, resampling settings and multiplicity adjustments are in the Supporting Information.

All three statistics correctly identified the identical scenario as non-significant and the location shift as significant. In the density-change scenario, SD and CSD detect the perturbation while ED does not at the 0.05 level in this example.

For a uniform contraction, the barycenter is preserved and the mean cross-distance shifts only modestly. SD detects the change because the contraction tightens the intra-distribution signature of the contracted set relative to the cross-signature, generating a systematic shape difference in the sorted arrays. Across repeated comparisons at this sample size, SD and CSD detect the contraction more reliably than ED. Rejection rates, uncertainty and additional sample sizes and comparators are reported in Supporting Figure S3.

### Per-point loss landscapes

Beyond aggregate distance computation, per-point penalties illustrate the geometric mechanisms of the ED cross-attraction component and the mean signature-profile comparison. For each candidate point *g* generated against a true data manifold *Y*, we define the pointwise generative penalties *δ*(*g*). For the cross-attraction component of Energy Distance, this evaluates the expected physical distance to the manifold: *δ*_ED_(*g*) = E_*y*∼*Y*_ ∥*g* − *y*∥. For the signature-profile illustration, this evaluates the radial-profile divergence of *g*’s local neighborhood: *δ*_SD_(*g*) = *W*_1_(*S*_*gY*_, 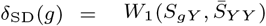 ), where *S*_*gY*_ = sort(∥*g* − *Y* ∥) is the cross-signature from *g* to *Y*, and 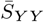 is the expected intra-manifold signature.

We evaluated these landscapes on two topologies that highlight the difference between these pointwise penalties: (a) two separated clusters and (b) a continuous ring (Figure 3).

**Fig. 3:**
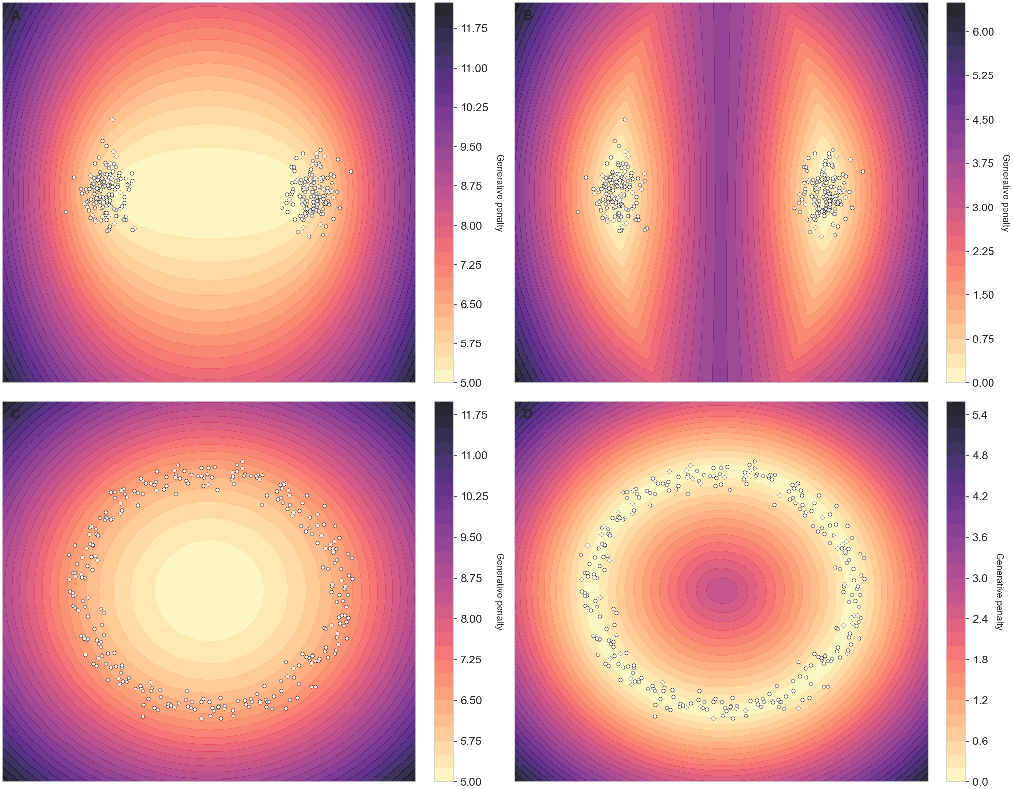
Per-point generative loss landscapes reveal how a single new point would be penalised at each location. *Top row (two clusters):* the cross-attraction landscape (left) is nearly flat in the inter-cluster gap, offering no gradient signal to guide a generated point away from the empty region; the mean-profile penalty (right) produces a pronounced penalty ridge between the clusters, showing that a point placed there would have an anomalous distance profile relative to both cluster distributions. *Bottom row (ring):* the cross-attraction minimum is at the ring’s empty centre (where mean distance to all reference points is minimised), while the mean-profile minimum traces the ring perimeter itself.

In the two-cluster scenario, a point *g* placed in the equidistant gap satisfies E∥*g* −*Y* ∥ ≈ E_*y*∼*Y*_ ∥*y* −*Y* ∥: the mean physical distance to the opposing cloud matches that of a genuine in-distribution point, so cross-attraction assigns a similar penalty to such a candidate as to real data, resulting in an uninformative landscape in the gap region. By contrast, the full radial profile of *g* differs from the mean within-reference profile 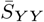, so *δ*_SD_(*g*) ≫ 0: the spurious candidate receives a large penalty that cleanly separates it from genuine data points.

The ring topology reveals a mismatch between the attraction minimum and the ring: for an ideal symmetric ring, its geometric center uniquely minimizes E∥*g* − *Y* ∥, by symmetry and convexity. This attraction minimum therefore lies in the empty interior. The SD penalty at the center is instead large: all ring points lie at approximately the same distance from the center, so *S*_*gY*_ is a nearly constant array (a flat profile), while the true intra-ring signature 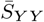 spans distances to both nearby and distant ring points. This shape mismatch produces *δ*_SD_(*g*) ≫ 0, and the SD minimum aligns with the ring perimeter, consistent with the contour plots in Figure 3.

### Interpolation sensitivity

Two TCGA sub-populations (*A* and *B*, 400 points each) were used to create interpolated points: *A*_*α*_ = (1−*α*)·*A*_matched_+*α*·*B*_matched_, where matching was performed via optimal assignment (Kuhn, 1955). SD(*A*_*α*_, *B*), CSD(*A*_*α*_, *B*), and ED(*A*_*α*_, *B*) were measured as *α* increased from 0 to 1. Each curve is divided by its starting value, *D*(*A, B*), so that all start at one and end at zero. The grey dashed line marks the common starting value of one.

ED decreased monotonically toward zero, while SD and CSD exhibited non-monotonic structural penalties, peaking near *α* = 0.5 (Figure 4). At *α* = 0.5, ED decreased to 0.81 of its starting value, while SD assigned an increased structural penalty of 1.45. CSD showed the strongest relative penalty (1.64), combining the row-level signature mismatch of SD with the column-level density mismatch of CD.

**Fig. 4:**
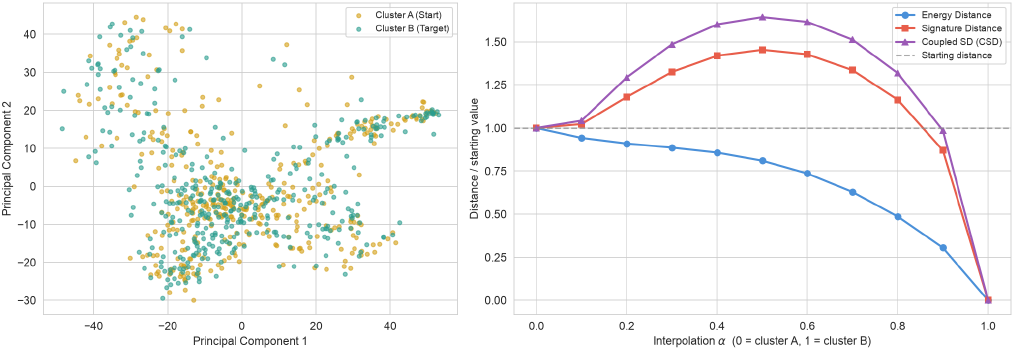
Interpolation sensitivity on TCGA transcriptomic data, testing whether distributional distances can detect synthetic samples that lie “between” real populations. *A* and *B* are 400-sample sub-populations; interpolated sets *A*_*α*_ = (1 − *α*)*A* + *αB* are constructed for *α* ∈ [0, 1]. *Left:* PCA projection of *A* (gold) and *B* (teal). *Right:* all three distances divided by their respective starting values; the grey dashed line marks the starting value of one. ED decreases monotonically as the interpolated set physically converges toward *B*, and never exceeds its starting value; it assigns no increased penalty to intermediate interpolants. SD peaks at *α* ≈ 0.5 and CSD shows an even stronger non-monotonic penalty, reflecting that the sorted distance profiles of interpolated points are systematically anomalous even when their mean-distance aggregate decreases. All distances are zero at *α* = 1, where *A*_*α*_ = *B*.

This divergence has a direct geometric explanation in terms of the signature shape argument developed in Section 3.2. Linearly interpolated points change the within- and cross-distribution distance profiles between *A* and *B*. SD registers this signature mismatch; ED, which collapses the distance profiles to their means, decreases along this interpolation. Supporting Figure S5 adds MMD and sliced *W*_1_ to this same experiment; Supporting Figure S4 gives a separate, independent-reference comparison over patient-disjoint partitions.

### Data expansion via guided Markovian sampling

Because SD is differentiable via automatic differentiation through `torch.sort`, the gradient ∇_*x*_SD(pop, *A*_*k*_) is computable using the subgradient conventions at ties and coincident points, making SD (and its extensions) usable directly as the potential energy of a Langevin update (Welling and Teh, 2011). This requires no generative model: the expansion operates as gradient descent in the space of empirical distributions, regularised by Cholesky-preconditioned Gaussian noise that confines displacement to directions of high variance in the seed set.

Using TCGA data clustered via Leiden (Traag et al., 2019), the largest cluster (931 patients) was partitioned into seed *A*, validation *B*, and hidden test *C* (310 each; one unused). Clustering, initialization and update parameters are specified in the Supporting Information. An initial population was constructed from *A* using *K*-means (Lloyd, 1982) centroids scaled to match |*A*|. The population was then evolved for 500 steps under Langevin dynamics:

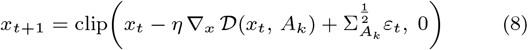

where 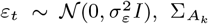 is the empirical covariance of the seed set (Cholesky-normalised), and *D* ∈ {ε, *S*, CSD} is the guidance loss. Critically, the gradient is taken with respect to the candidate population, using *A*_*k*_ only as the training reference: *B* is deliberately excluded from the training signal to preserve it as an uncontaminated validation reference.

The chain is stopped at the epoch that minimises the following stopping score, using *D* = *d*_*E*_ for ED guidance and *D* = SD for SD/CSD guidance:

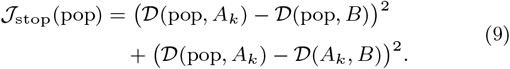

This objective is satisfied when *D*(pop, *A*_*k*_) ≈ *D*(pop, *B*) ≈ *D*(*A*_*k*_, *B*): the expanded population has become equidistant from *A* and *B* in the chosen discrepancy, under the stopping score.

To stabilise the stopping epoch, we applied a bootstrap protocol: *K* = 5 replicates draw *A*_*k*_ with replacement from *A*, each producing an independent chain with its own covariance 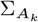. All five replicates are evaluated, and the replicate with the lowest validation distance at its selected stopping step is displayed; *C* is used exclusively for final evaluation and never influences stopping.

Both SD- and ED-guided traces show clear U-shaped minima within 500 steps, as does CSD-guided expansion (Figure 5). The SD-guided stopping epochs are stable across bootstraps (218 ± 12 steps), while ED-guided epochs show higher variance (174 ± 72 steps; mean ± bootstrap standard deviation).

**Fig. 5:**
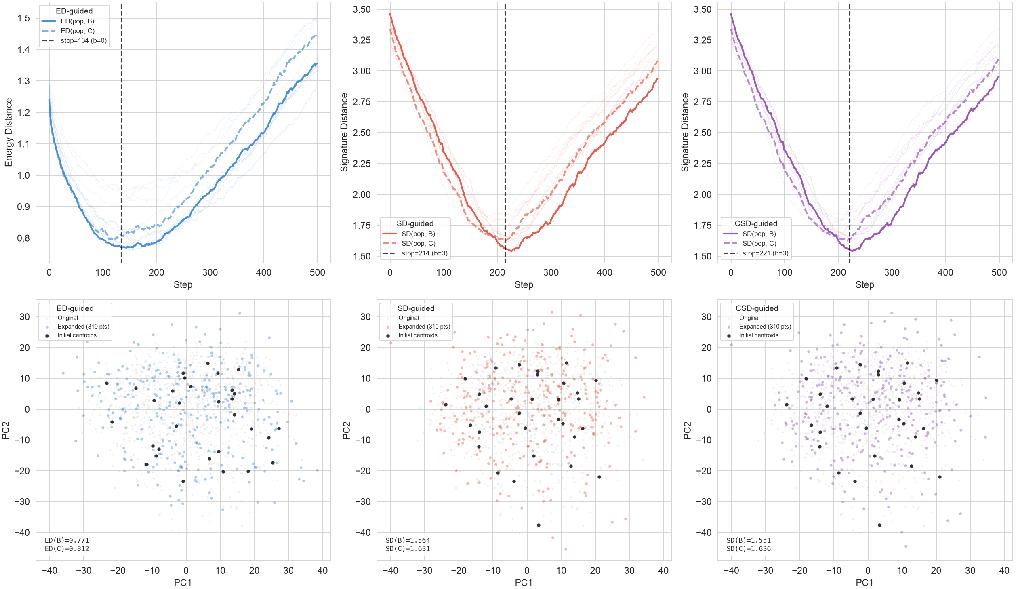
Langevin expansion on the largest TCGA Leiden cluster, demonstrating SD as a potential energy for model-free data augmentation. New samples are generated by gradient descent on the distributional distance to the cluster, then stopped when the validation stopping score *J*_stop_ (Eq. 9) reaches its minimum. *Top row:* distance traces across 500 steps for the validation-selected replicate among *K* = 5 bootstrap replicates; solid line = distance to validation set *B*, dashed = distance to held-out *C*; light traces show all bootstrap replicates; vertical dashed line marks the stopping epoch. All three objectives guide expansion; independent precision, recall and sliced Wasserstein are reported for all replicates in the Supporting Information. *Bottom row:* PCA projections at the stopping epoch; grey = original cluster, coloured = expanded samples, black dots = initial *K*-means centroids. Metrics annotated bottom-left.

The stopping criterion (Equation 9) encodes a soft indistinguishability constraint: the expanded population should be as similar to *B* as the seed *A*_*k*_ is, under the chosen discrepancy. The resulting U-shaped stopping score provides a principled, observable criterion whose minimum can be identified from the validation fold alone. The held-out fold *C* is used exclusively to verify that the stopping generalises; the gap between D(pop, *B*) and D(pop, *C*) at the selected step provides a held-out check of the stopping epoch.

Because the expansion uses only a local reference set to anchor the generated points in feature space, it can be applied independently to each subpopulation in a dataset. Partitioning the data and expanding each partition separately allows each observed subpopulation to be expanded, with the reference serving only to pin each expansion to the correct region of the space.

### SD as differentiable generative loss

Because the quantile sorting operations underlying the signatures are tracked by automatic differentiation frameworks (e.g., `torch.sort`), SD can be used as a differentiable loss function for training generative neural networks (implementation details in Appendix B). We first assess gradient-based training on controlled synthetic problems, then demonstrate the approach at scale on a tissue-conditioned generation task using TCGA gene expression data.

A class-conditioned multilayer perceptron (MLP) generator (five hidden layers with rectified linear unit, ReLU, activations) was trained to map isotropic Gaussian noise to a target distribution of two intertwining circles of different radii (Figure 6). All models were evaluated by SD and ED score (lower is better) on 2,000 generated versus 2,000 independent real points per class. Three independent training runs are evaluated per loss; a prespecified run is displayed. Repeated results, Gaussian MMD and sliced *W*_1_ loss comparisons, and independent precision/recall estimates are in Supporting Figures S6–S7 and the supporting tables.

**Fig. 6:**
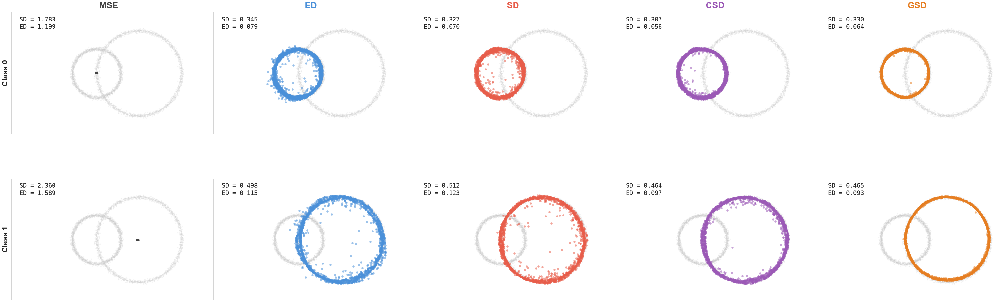
Generative loss comparison on two intertwining circles of different radii, a geometry that the mean squared error (MSE) loss does not recover in this unpaired task. Each column shows a different loss function; rows show class 0 (inner circle) and class 1 (outer circle). MSE collapses both classes to their respective barycenters, producing point masses. All four distributional losses recover the circular structure, confirming that distributional objectives can learn topology from an unpaired generation task where MSE cannot.

MSE collapses to the class-conditional barycenter, as expected when a pointwise loss is applied to an unpaired generation task. The distributional losses all recover the circular structure. We additionally tested network normalization and loss decomposition on the same task (Figure 9, Appendix B). In this experiment, batch normalization causes fragmentation of the generated distribution (Appendix B).

#### Heteroscedastic sinusoidal generation

To illustrate that these properties extend from topology-learning to continuous distribution learning, we trained a noise-conditioned MLP with two hidden layers on a heteroscedastic sinusoidal task: *y* = sin(*t*)+ *σ*(*t*) · *ξ*, with narrow noise (*σ* = 0.06) for *t* ∈ [0, *π*] and wide noise (*σ* = 0.28) for *t* ∈ [*π*, 2*π*] (Figure 7). The model takes (*t, ε*) as input and generates both coordinates (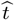, *ŷ*). The noise *ε* is never paired with the specific *ξ* used to draw a target, so a point-prediction loss has little incentive to exploit it; distributional losses, by contrast, naturally use *ε* to span the output distribution.

**Fig. 7:**
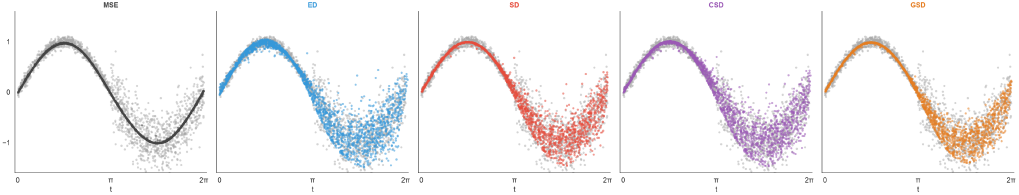
Heteroscedastic sinusoidal generation (*y* = sin(*t*) + *σ*(*t*)*ξ*, narrow noise for *t* ∈ [0, *π*], wide noise for *t* ∈ [*π*, 2*π*]). The noise input *ε* is never paired with the specific *ξ* used to draw the target, so the model must learn to use *ε* to span the output distribution. MSE collapses to the conditional mean, discarding the noise input entirely. Distributional losses recover different amounts of heteroscedastic spread. Three independent runs are evaluated and a prespecified run is displayed; repeated output-spread diagnostics are reported in the Supporting Information.

The distributional losses recover heteroscedastic structure, with different amounts of output spread. Joint and residual diagnostics across seeds are in the Supporting Information.

#### Tissue-conditioned generation on TCGA landmark genes

Having assessed gradient-based training on controlled tasks, we applied the losses to a biologically relevant generation problem designed to expose how each loss function handles high-dimensional, multi-population structure under a deliberate capacity constraint. TCGA provides RNA expression measurements across 978 landmark genes (Subramanian et al., 2017) and 24 supplied cohort-derived tissue labels. A tissue-conditioned generator was trained to map a low-dimensional noise vector (*ε* ∈ ℝ^32^) concatenated with a tissue-type label to a 978-dimensional gene expression profile. The 32-dimensional input is a deliberate bottleneck: with 978 output dimensions, a higher-capacity input could reduce the performance gap between losses, masking their technical differences. The constrained regime forces each loss to prioritise different aspects of the distributional geometry, revealing their distinct inductive biases.

Each loss was trained with a *glocal* protocol, combining a global term (the distance computed on the full mixed-tissue batch) with local per-tissue terms (the mean of per-tissue distances within each tissue group in the batch):

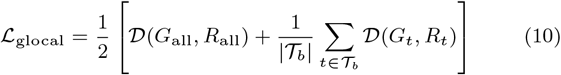

where *T*_*b*_ is the set of tissue labels represented by at least two observations in the batch, and *G*_*t*_, *R*_*t*_ are the generated and real samples for tissue *t*. The global term targets cross-tissue manifold structure; the local terms target within-tissue distributional fidelity. All losses are trained with the same protocol, using five seeds each; ED and SD optimize 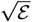 and 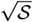, respectively. The data comprise 5,548 fitting, 2,378 validation and 1,994 test observations, partitioned by patient, with normalization fitted on the fitting data. The full sample manifest, preprocessing and training settings are in the Supporting Information.

The quality of each generator was evaluated on a patient-disjoint held-out test set of 1,994 samples via two independent criteria: (1) a downstream tissue-type classifier trained exclusively on 10× synthetic data and evaluated on real test samples (accuracy), measuring whether the synthetic data captures tissue-discriminative structure; and (2) distributional recall on held-out real samples, using *k* = 3 nearest-neighbour balls in the full feature space.

The glocal protocol reveals different strengths among distributional losses (Figure 8). All losses achieve useful classification accuracy (85.6–92.7%), including MSE, whose local term learns tissue-specific means from within-tissue batch pairings. SD and CSD achieve higher mean distributional recall (19.2% and 20.2%, respectively, versus 12.6% for ED), recovering more of the held-out distribution. Per-tissue F1 scores (Figure 10, Appendix B) show how performance varies across tissues. Macro-F1 and independent precision/recall comparisons are in the Supporting Information.

**Fig. 8:**
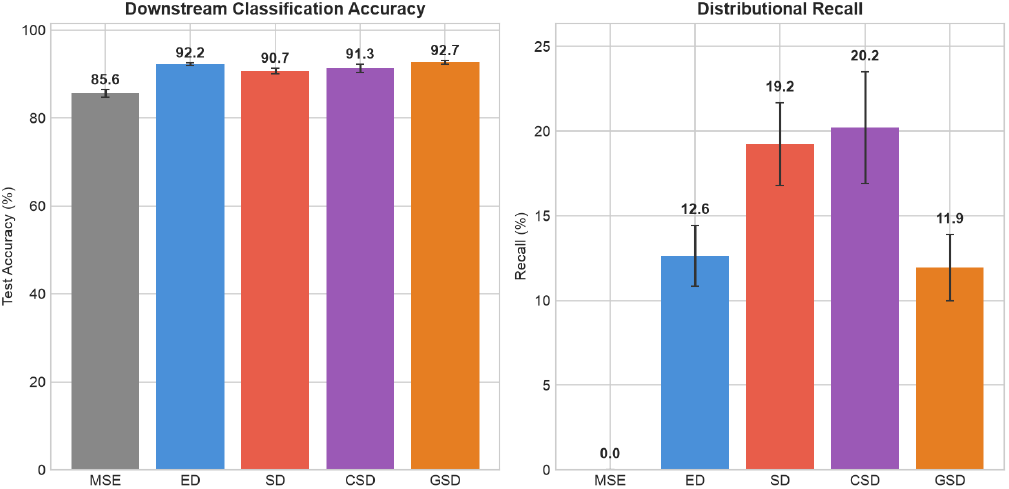
Downstream tissue classification accuracy (left) and distributional recall (right) on real test data. Classifiers are trained exclusively on synthetic data. SD and CSD recover more of the held-out distribution, while retaining strong classification accuracy. Bars show means and standard deviations across five seeds.

## Discussion

### Interpretation and broader implications

The sensitivity and loss landscape experiments share a common geometric explanation. Energy distance collapses the full distance profile from a point to the opposing distribution into its mean, discarding the rest of each individual profile. Signature Distance retains the full sorted profile, making it sensitive to any distributional property encoded in the shape of the radial distance distribution: variance, skewness, and multi-modality. The advantage of SD over ED is therefore determined by whether the distributional difference in question alters the shape of these profiles. The signature matrix encodes complementary information along its two axes: CD captures population-level density-scale constraints by sorting each column independently, while SD captures per-point topology via rows. Their Pythagorean combination (CSD) adds within-population density constraints, while the nearest-neighbour grounding in GSD addresses this more directly, preserving point identity at the same asymptotic cost.

A practical consequence for generative model evaluation in biology (Lopez et al., 2018; Marouf et al., 2020) is that models suffering from interpolation artifacts (Goodfellow et al., 2014; Kingma and Welling, 2014; Rezende and Mohamed, 2015) can produce synthetic samples that receive favorable scores under evaluators such as ED and kernel MMD despite lacking realistic local structure. SD provides a more sensitive criterion for the interpolation artifacts illustrated here because it evaluates the full distance profile shape rather than its mean.

The model-free character of the Langevin expansion is particularly relevant for biological data augmentation: no generative architecture is required, the noise covariance is estimated directly from the seed cluster, and distributional distances guide both the objective and evaluation. Unlike interpolation-based augmentation methods (Chawla et al., 2002), which can produce interpolation artifacts penalized by SD (Section 3.3), the Langevin approach generates samples by matching the distributional geometry of the seed population. This makes it applicable to any modality for which a pairwise distance can be computed, such as protein structures, chemical fingerprints or multi-omic profiles (Argelaguet et al., 2018), without any retraining. It is especially relevant in settings where foundation model embeddings are available but task-specific labels are not. In single-cell biology, for example, pretrained encoders (Theodoris et al., 2023; Cui et al., 2024) provide rich representations of cellular state, yet labelled examples of a specific perturbation effect (Dixit et al., 2016) may comprise only a handful of observations. Distributional losses allow new samples to be generated directly in the embedding space, guided solely by the geometry of the observed population, without requiring paired supervision or finetuning of the foundation model.

Distributional losses address a different problem from pointwise regression: they match *distributions* where no pointwise correspondence exists. They serve as natural objectives for unconditional generation, data augmentation, and model-free sampling, and may complement pointwise losses as hybrid objectives or as alternatives to adversarial discriminators where mode coverage is desirable. The glocal protocol (Equation 10) illustrates the role of batch composition on multi-population data, but the design space of batch strategies remains largely unexplored.

Density-peaks clustering (Rodriguez and Laio, 2014) also uses local density, but selects centres and partitions observations; SD compares two radial-distance distributions without requiring a cluster count.

### Limitations and future directions

SD requires sorting the distance profiles, adding O(*n*^2^ log *n*) work beyond the quadratic pairwise-distance computation. At larger scales, subsampling or approximate nearest-neighbor methods can reduce this cost, with a corresponding approximation of the statistic.

Further theoretical properties and related metrics are discussed in Appendix C. The hard nearest-neighbour assignment in GSD is a simple, parameter-free profile-matching rule that does not impose transport mass constraints; smoother alternatives, such as softmax-weighted profile matching or Sinkhorn-based optimal plans (Cuturi, 2013) over signature rows, may improve calibration at the cost of additional computation.

SD is defined for any metric space, requiring only pairwise distances, making it applicable to protein structure comparison (RMSD-based distances), chemical similarity (Tanimoto distances), and multi-omic integration. The distance signature shares underpinnings with the Vietoris–Rips filtration in computational topology (Edelsbrunner and Harer, 2010): a point’s signature encodes the birth times of its incident edges in a Vietoris–Rips complex. Formalizing connections to persistent homology may yield further theoretical insights.

### Concluding remarks

We introduced Signature Distance, a distributional comparison method that generalises energy distance by retaining the full sorted pointwise distance profile rather than collapsing it to a scalar mean. SD matches ED’s quadratic pairwise-distance cost, with an additional sorting step, while capturing local density structure and interpolation artifacts through the full radial profiles. Extensions along complementary axes of the signature matrix (Column Distance for population-level density constraints, and nearest-neighbour grounding (GSD) for spatial correspondence) address different aspects of distributional fidelity.

On TCGA gene expression data, SD and its extensions translate these geometric properties into practical advantages: as two-sample tests, as differentiable generative training objectives with a glocal protocol for multi-population data, and as direct potential energies for model-free Langevin data expansion. The tissue-conditioned generation experiment demonstrates that under capacity constraints, distributional losses extract more useful structure for tissue classification than pointwise regression at the tested latent dimension, and that batch composition is a key design consideration when deploying these losses at scale. Code and data to reproduce all experiments are available on GitHub.

## Data Availability

The TCGA data used in this work was downloaded from the GDC Data Portal (https://portal.gdc.cancer.gov/). Preprocessing and patient partitions are documented in the Supporting Information. The code used to perform the experiments and reproduce the results in this paper is available under the MIT license at https://github.com/lazzaronico/signature-distance/releases/tag/v0.2.1 and archived on Zenodo (doi:10.5281/zenodo.22767584). The repository reproduction guide provides the experiment configurations, preprocessing scripts and individual-run results.

## Acknowledgments and Funding

The results shown here are in whole or part based upon data generated by the TCGA Research Network^1^. This work is partially funded by the EU through the 3DSecret project under the HORIZON-EIC-2022-PATHFINDER-OPEN-01-01 programme (grant no. 101099066).

## Appendix A Upper and Lower Bounds for Squared Signature Distance

### Theorem 1.

*Let µ and ν be probability measures on* ℝ^*d*^ *with finite first moments. Define the population-level asymmetric divergence*

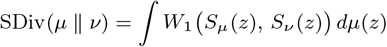

*where S*_*µ*_(*z*) *denotes the distribution of radial distances from z under µ (i*.*e. the pushforward* (*f*_*z*_)#*µ with f*_*z*_(*u*) = ∥*z* − *u*∥*), and the symmetric squared distance S*(*µ, ν*) = SDiv(*µ* ∥ *ν*) + SDiv(*ν* ∥ *µ*). *Then:*

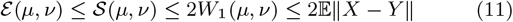

*Proof of the upper bound (*S ≤ 2*W*_1_). For any reference point *z* ∈ ℝ^*d*^, the signature *S*_*µ*_(*z*) = (*f*_*z*_)#*µ* is the pushforward of *µ* through *f*_*z*_(*u*) = ∥*z* − *u*∥. By the reverse triangle inequality, *f*_*z*_ is 1-Lipschitz: |*f*_*z*_(*u*) − *f*_*z*_(*v*)| = ∥*z* − *u*∥ − ∥*z* − *v*∥ ≤ ∥*u* − *v*∥. A standard property of optimal transport states that the 1-Wasserstein distance is non-increasing under 1-Lipschitz pushforwards: any coupling *γ* for (*µ, ν*) induces a coupling for (*S*_*µ*_(*z*), *S*_*ν*_ (*z*)) with cost |*f*_*z*_ (*u*)−*f*_*z*_ (*v*)| *dγ* ≤ ∥*u*−*v*∥ *dγ*; taking the infimum gives

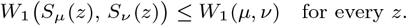

Taking the expectation over *z* ∼ *µ* establishes SDiv(*µ* ∥ *ν*) ≤ *W*_1_(*µ, ν*); by symmetry, SDiv(*ν* ∥ *µ*) ≤ *W*_1_(*µ, ν*); their sum S is bounded by 2*W*_1_. Finally, *W*_1_(*µ, ν*) ≤ E ∥*x* − *y*∥ because the independent coupling is a valid (non-optimal) transport plan.

*Proof of the lower bound (*S ≥ ε ). For any two one-dimensional distributions *A* and *B*, the 1-Wasserstein distance satisfies the mean bound: *W*_1_(*A, B*) ≥ | E [*A*] − E [*B*]| (the identity function is 1-Lipschitz, so this follows from the Kantorovich–Rubinstein duality with test function *f* (*t*) = *t*). Applying this to the recall term at reference *y* ∼ *ν*:

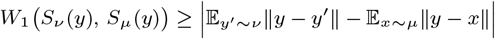

Averaging over *y* ∼ *ν* and applying Jensen’s inequality (E |*h*| ≥ |E [*h*]|):

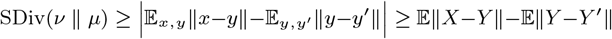

where the last step uses |*a* − *b*| ≥ *a* − *b*. By exact symmetry, SDiv(*µ* ∥ *ν*) ≥ E ∥*X* − *Y* ∥ − E∥*X* − *X*^′^∥. Summing the two lower bounds and collecting:

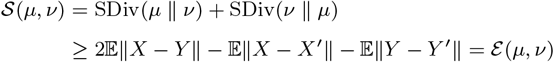

When E∥*X* − *Y* ∥ = 0, normalized SD is defined as zero. The energy lower bound implies separation on Euclidean probability measures with finite first moments. Symmetry and nonnegativity follow by construction. Appendix C establishes value stability under Wasserstein perturbations.

## Appendix B Signature Distance as a Differentiable Generative Loss

Because autograd tracks indexing operations during torch.sort, SD^2^ is piecewise differentiable and can be used as a structural loss function for deep generative modeling. The following implementation choices are recommended for training with distributional losses. Figure 9 reports exploratory component comparisons; their estimator and the training protocols are specified in Section S3. The synthetic and global-only comparisons optimize raw ε and S; the glocal TCGA comparison optimizes ED and SD.

**Fig. 9:**
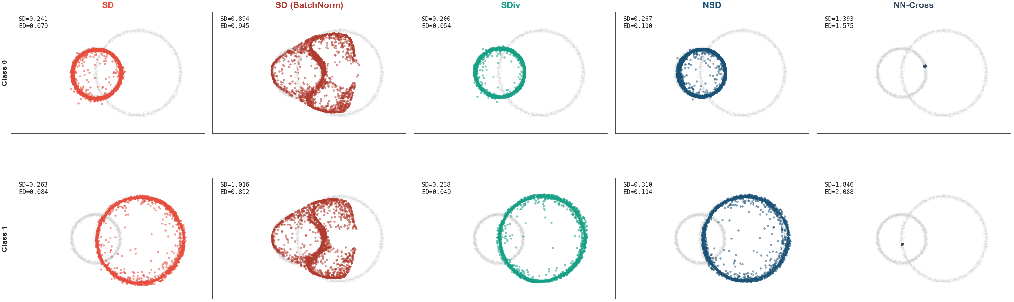
Normalization and component analysis on the ring generation task (class 0 on top, class 1 on bottom). *Left pair:* SD succeeds without normalization but fails catastrophically with BatchNorm. BatchNorm normalises activations *across the batch*, changing the batch geometry in this run: the generated distribution fragments into disconnected clusters rather than forming a coherent ring. *Right panels:* component ablation. The recall term SDiv alone (with cached real profiles) recovers the ring, while the NN-Cross term alone (without profile comparison) collapses entirely, illustrating the effect of including profile matching in these runs.

**Fig. 10:**
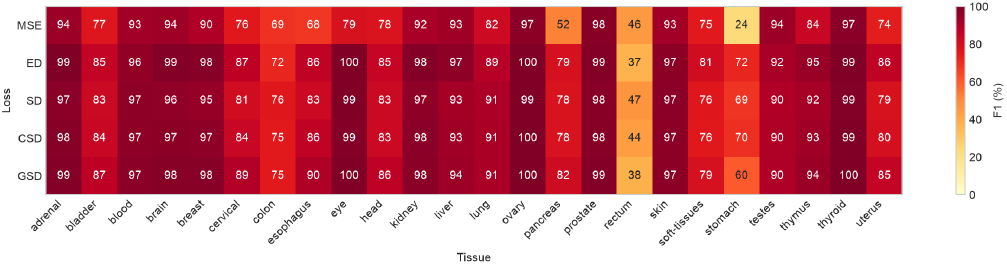
Per-tissue F1 scores for the tissue-conditioned generation experiment (noise dim = 32, glocal protocol). Each cell shows the F1 score (%) for a given tissue–loss combination, with a classifier trained exclusively on synthetic data and evaluated on real test samples. Cells show means across five training seeds on patient-disjoint partitions. Label-level performance varies; the small stomach-labelled group warrants caution.

### Normalization

BatchNorm can affect distributional training: it enforces zero-mean unit-variance per activation within the batch, and can alter the batch geometry used by a distributional objective. Figure 9 demonstrates this on the ring task: SD with BatchNorm produces visibly fragmented, mode-collapsed outputs, while the same loss without normalization recovers the target distribution. The figure also shows three additional losses not in the main comparison: SDiv (recall-only) and NSD both recover the ring structure, while the NN-Cross term alone fails completely, suggesting a role for the profile comparison backbone in this example. LayerNorm (normalizing per sample, not per batch) is an alternative. No normalization at all is also viable when inputs are pre-normalized to unit variance.

### Gradient flow

Detaching the within-set signature profiles from autograd (treating *S*_*XX*_ and *S*_*Y Y*_ as zero-gradient constants) degraded performance in the exploratory runs. The gradient flowing through *S*_*XX*_ contributes a density-conditioning signal.

### Global-only comparison

Results from the same architecture and patient partitions, trained without per-tissue terms, are reported in Section S3. These runs optimize raw energy and signature objectives with stochastic validation.

## Appendix C Measure-Theoretic Formulation

The discrete algorithm in Section 2.1 is the natural estimator of a continuous population-level quantity. Let *µ* and *ν* be probability measures on ℝ^*d*^ with finite first moments. For any reference point *z*, the sorted signature represents the empirical radial-distance distribution; in the population limit, at radius *r* this becomes the CDF *F*_*zµ*_(*r*), equal to the measure of the closed ball *B*_*r*_(*z*):

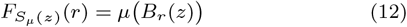

Because the *W*_1_ distance between two one-dimensional distributions equals the *L*_1_ integral of the absolute difference of their CDFs, the pointwise divergence at reference *z* is:

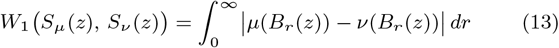

Substituting into the symmetrized aggregate over all reference points, with the sum measure *µ* + *ν*, yields the continuous form of Squared Signature Distance:

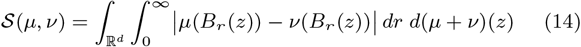

This formulation makes precise the sense in which SD is a local projection of the optimal transport geometry: it measures how much the two distributions differ in their ball-mass assignments, integrated over all radii and reference locations.

For the first transport identity below, every point coupling pushes forward to a coupling of the radial laws. Conversely, conditional distributions of points given their radii lift a radial coupling to a point coupling with the same radial cost. Such conditional distributions exist on Euclidean space. This proves equality of the infima.

### Bridge to full Wasserstein distance

An explicit bridge to full transport uses the same radial observations. Put *g*_*z*_(*x, y*) = |∥*x* − *z*∥ − ∥*y* − *z*∥| and let Π(*µ, ν*) denote all couplings. Then

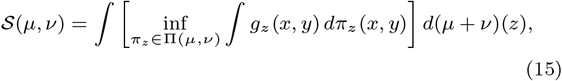

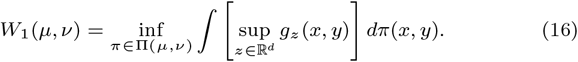

The supremum equals ∥*x*−*y*∥ by the reverse triangle inequality and equality at *z* = *x*. For empirical inputs, taking the maximum over *X* ∪ *Y* is already exact. Thus SD compares radial marginals using separate anchor-specific matchings and a sum of anchor averages; full *W*_1_ retains one common point matching and uses a maximum radial discrepancy for each matched pair.

### Fixed-reference metric

Put *h*_*z*_(*µ, ν*) = *W*_1_(*S*_*µ*_(*z*), *S*_*ν*_ (*z*)). For one fixed probability measure *Q* used for every pair, define

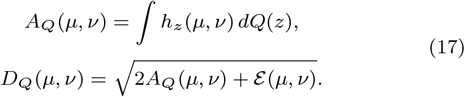

Anchor-wise triangle integrated against the common *Q* proves that *A*_*Q*_ is a pseudometric. If *Q* has full support in ℝ^*d*^, continuity of *h*_*z*_ implies that *A*_*Q*_ = 0 forces *h*_*z*_ = 0 everywhere, hence S = 0 and equality of measures: *A*_*Q*_ is then already a metric. The square root of *A*_*Q*_ is always a pseudometric. Applying the Euclidean Minkowski inequality to the pair 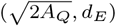 proves triangle for *D*_*Q*_, and energy supplies separation.

### Perturbation stability

For another pair *µ*^′^, *ν*^′^ with finite first moments, put *h*_*z*_ = *W*_1_(*S*_*µ*_(*z*), *S*_*ν*_ (*z*)) and *δ*= *W*_1_(*µ, µ*^′^) + *W*_1_(*ν, ν*^′^). Then

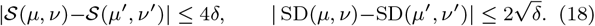

To see this, radial contraction and the reverse triangle inequality for one-dimensional *W*_1_ give |*h*_*z*_(*µ, ν*) − *h*_*z*_(*µ*^′^, *ν*^′^)| ≤ *δ*uniformly in *z*. Also, for any fixed pair, *z* 1→ *h*_*z*_ is 2-Lipschitz: moving the anchor by ∥*z* − *w*∥ changes each radial law by at most that amount in *W*_1_. First changing the compared measures while retaining the anchor measure *µ*+*ν* therefore costs at most 2*δ*; then changing that anchor measure to *µ*^′^ +*ν*^′^ costs at most another 2*δ*, by coupling the anchors. The square-root bound follows from 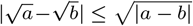 for nonnegative *a, b*.

### Discrete computation

For a reference point *z*, sort the distances to *X* and *Y* into *a*_(1)_(*z*) ≤ · · · ≤ *a*_(*n*)_(*z*) and *b*_(1)_(*z*) ≤ · · · ≤ *b*_(*m*)_(*z*). For 0 *< u <* 1, their empirical quantiles are *q*_*X*_ (*z, u*) = *a*_(⌈*nu*⌉)_(*z*) and *q*_*Y*_ (*z, u*) = *b*_(⌈*mu*⌉)_(*z*). All observations retain their empirical mass, including zero self-distances.

When *n* = *m* = *N*, the integral is the mean absolute sorted-rank difference:

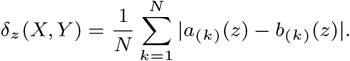

For unequal sizes, let 0 = *t*_0_ *< t*_1_ *<* · · · *< t*_*L*_ = 1 be the ordered union of the breakpoints 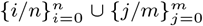, and set *u*_*l*_ = (*t*_*l*−1_ + *t*_*l*_)*/*2. Exact integration gives

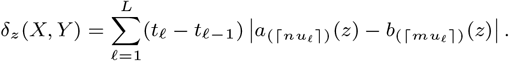

The empirical expectations in the main definition are therefore

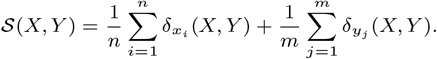

In matrix notation, *D*_*XX*_ [*i, j*] = ∥*x*_*i*_ − *x*_*j*_ ∥, *D*_*XY*_ [*i, j*] = ∥*x*_*i*_ − *y*_*j*_ ∥, and *D*_*Y Y*_ [*i, j*] = ∥*y*_*i*_ − *y*_*j*_ ∥. The intra-signature *S*_*XX*_ (*x*_*i*_) and cross-signature *S*_*XY*_ (*x*_*i*_) are the sorted rows of *D*_*XX*_ and *D*_*XY*_ ; the corresponding real-reference signatures are sorted rows of *D*_*Y Y*_ and 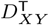 .

## Appendix D Analytic Reduction to Energy Distance

Signature distance is a structural generalization of energy distance (Székely and Rizzo, 2013), whose raw expression is:

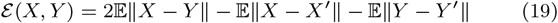

With Δ_*z*_(*u*) = *q*_*ν*_ (*z, u*) − *q*_*µ*_(*z, u*), the first-moment identity gives

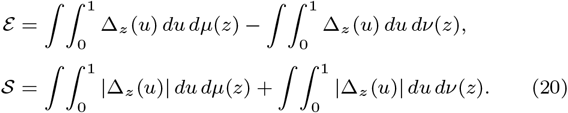

The signed terms of energy are replaced by absolute radial-quantile differences in *S*. Equality holds if and only if Δ_*z*_ ≥ 0 at *µ* anchors and Δ_*z*_ ≤ 0 at *ν* anchors, almost everywhere in anchor and quantile: the two contributions to *S* − ε are integrals of the nonnegative functions |Δ| − Δ and |Δ| + Δ, respectively. For bounded supports, it is sufficient that every cross-distance exceed both within-support diameters. Equality need not require that strong separation condition. Conversely, overlap alone does not imply strict inequality. Energy remains a separating distributional statistic even though it uses first moments of distance profiles; retaining the full profile changes finite-sample sensitivity, not the existence of a population distinction.

## Appendix E Additional Experiments and Methods

### S1. Datasets and preprocessing

The landmark input contains 978 genes and 24 cohort-derived labels (Subramanian et al., 2017).

**Fig. S1:**
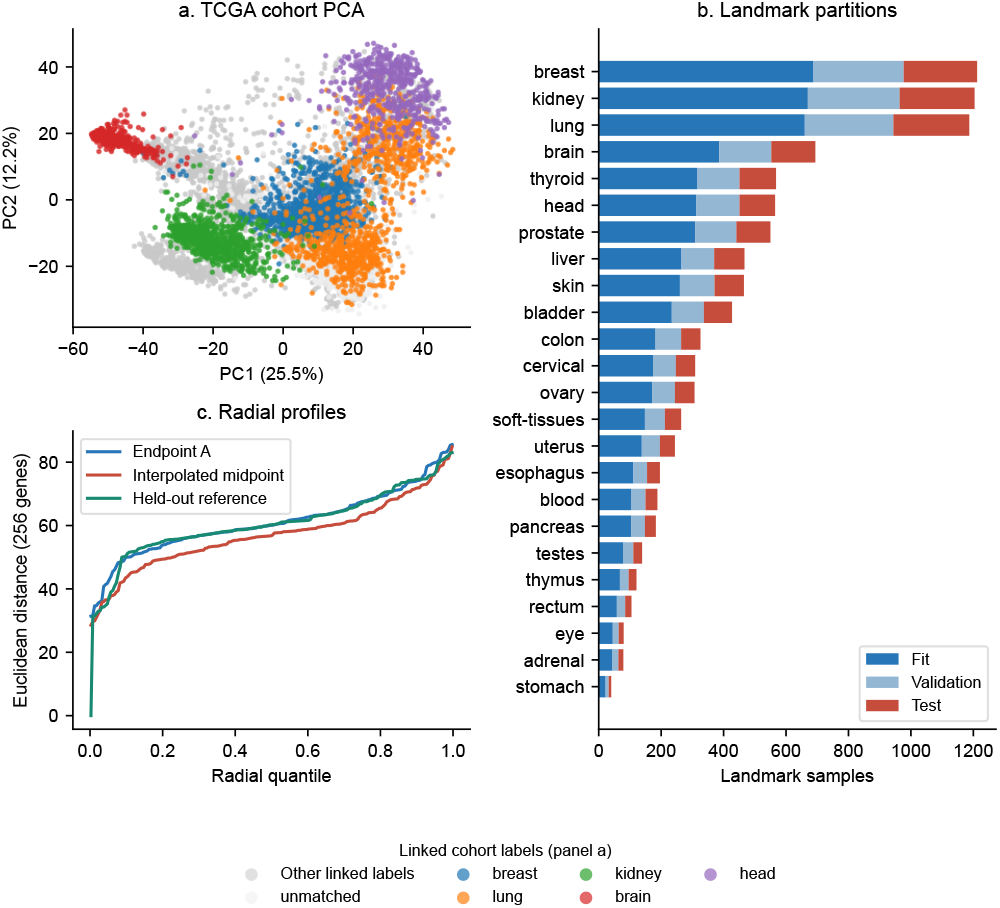
(a) Upper left: PCA of the 8,954-patient general RNA-seq representation, showing the five most numerous linked label groups separately; other linked labels and unassigned observations are shown in grey. The first two PCs explain 25.5% and 12.2% of variance. Numerical distances use all 256 selected expression features. (b) Right: class counts in the separately processed 978-gene landmark task, divided by patient into fitting, validation and test. (c) Lower left: radial quantiles around an illustrative held-out reference point, showing construction endpoint A, interpolated midpoint and held-out reference.

#### Landmark sample counts

| Label | Fit | Validation | Test |
| --- | --- | --- | --- |
| adrenal | 44 | 19 | 16 |
| bladder | 234 | 103 | 90 |
| blood | 105 | 45 | 38 |
| brain | 386 | 167 | 141 |
| breast | 688 | 289 | 235 |
| cervical | 174 | 73 | 62 |
| colon | 182 | 82 | 62 |
| esophagus | 111 | 44 | 41 |
| eye | 45 | 19 | 16 |
| head | 313 | 138 | 114 |
| kidney | 670 | 294 | 240 |
| liver | 265 | 105 | 97 |
| lung | 660 | 284 | 243 |
| ovary | 171 | 73 | 63 |
| pancreas | 104 | 44 | 35 |
| prostate | 309 | 132 | 109 |
| rectum | 58 | 27 | 20 |
| skin | 260 | 111 | 94 |
| soft-tissues | 148 | 64 | 52 |
| stomach | 22 | 10 | 8 |
| testes | 78 | 33 | 28 |
| thymus | 68 | 28 | 25 |
| thyroid | 315 | 136 | 117 |
| uterus | 138 | 58 | 48 |

The cohort-derived task labels group PCPG under kidney and MESO under lung.

After deduplication, the landmark cohort contains 9,920 aliquots from 9,202 patients. Identical-expression duplicate aliquots are retained once, preferring the specific ESCA annotation over the aggregate STES annotation where they conflict.

Patients, stratified by these labels, are divided into fitting, validation and test partitions containing 5,548, 2,378 and 1,994 aliquots from 5,152, 2,209 and 1,841 patients, respectively. No patient crosses partitions. Per-gene means and sample standard deviations are estimated on the fitting partition and applied to validation and test data.

### S2. Statistical comparisons

#### Permutation tests and sensitivity

The controlled comparison uses n=m in {50, 200}, five prespecified Gaussian scenarios, and 50 independently generated sample pairs per scenario and size (500 pairs in total). X follows N(0,I_2_), and independent Y follows N((a,0),s^2^I_2_), with (a,s) equal to (0,1), (2,1), (0,0.7), (0.25,1), or (0,0.9). All five statistics—E, S, CSD, Gaussian MMD^2^ and sliced W_1_—use common data and 999 pooled label permutations. The Gaussian-kernel bandwidth is the pooled median positive pairwise distance, fixed under permutations; sliced W_1_ uses 128 fixed normalized Gaussian directions. Rejection fractions are per-test rates at 0.05, with 95% Wilson intervals over the 50 independent pairs. Holm adjustment addresses the five simultaneous method comparisons of each illustrative pair separately. It is not applied to the per-method rejection fractions.

#### Raw and adjusted illustrative p-values

| Scenario | Statistic | P | Holm |
| --- | --- | --- | --- |
| Same Gaussian | E | 0.816 | 1.000 |
| Same Gaussian | S | 0.883 | 1.000 |
| Same Gaussian | CSD | 0.870 | 1.000 |
| Same Gaussian | MMD<sup>2</sup> | 0.804 | 1.000 |
| Same Gaussian | Sliced $W_1$ | 0.803 | 1.000 |
| Shift = 2 | E | 0.001 | 0.005 |
| Shift = 2 | S | 0.001 | 0.005 |
| Shift = 2 | CSD | 0.001 | 0.005 |
| Shift = 2 | MMD<sup>2</sup> | 0.001 | 0.005 |
| Shift = 2 | Sliced $W_1$ | 0.001 | 0.005 |
| Scale = 0.7 | E | 0.053 | 0.066 |
| Scale = 0.7 | S | 0.001 | 0.005 |
| Scale = 0.7 | CSD | 0.001 | 0.005 |
| Scale = 0.7 | MMD<sup>2</sup> | 0.009 | 0.027 |
| Scale = 0.7 | Sliced $W_1$ | 0.033 | 0.066 |

**Fig. S2:**
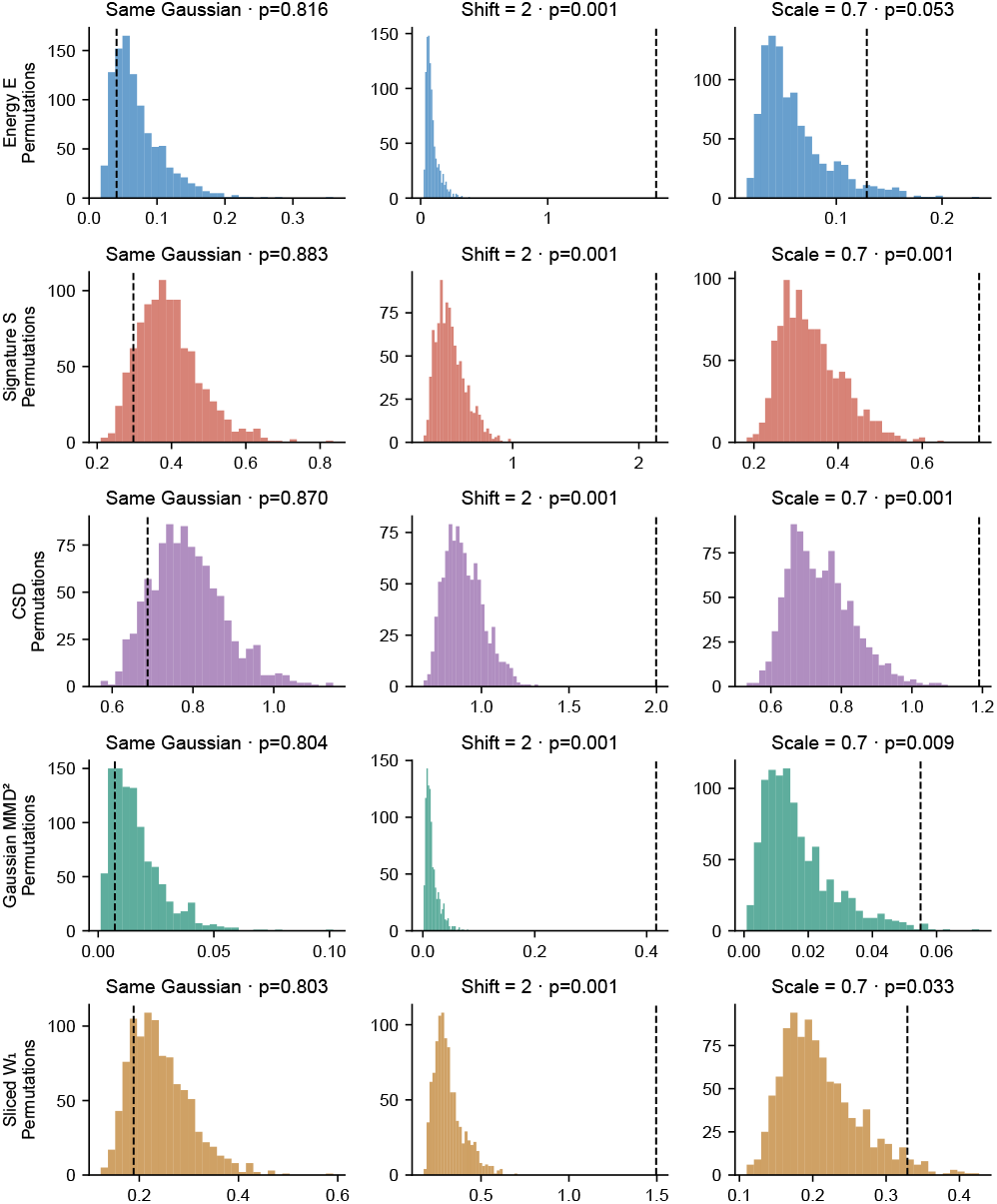
The first independent sample pair at n=50 for the null, large location shift and 0.7-scale contraction. Histograms show 999 pooled, size-preserving label permutations; dashed lines are the observed values. Displayed p-values are unadjusted plus-one values, with minimum 0.001. The table also gives Holm adjustment across the five statistics for each single sample pair. All 50 repetitions, including the n=200 comparison, appear in Supporting Figure S3.

#### Self-bootstrap references

Main Figure 1 uses two 30-point groups. The two directional panels use 500 self-bootstrap comparisons: for each reference Z (X or Y), draw two independent size-30 samples with replacement from Z and compute SDiv between them. Each reference distribution depends only on its source sample and resampling settings; it can be computed once and reused when the opposite target changes. The annotated tail fraction is (1 + number of self-bootstrap values at least as large as the observed cross-sample divergence)/501. It is a descriptive bootstrap-reference exceedance, not a finite-sample calibrated permutation p-value. The third, symmetric SD panel uses 999 size-preserving pooled label permutations. The grey bands in all three lower panels are separate 95% percentile bootstrap intervals from 500 independent within-group resamples. Each resample recomputes both directed statistics and their summed square root.

### S3. Generative applications

#### Shared training settings

We compare MSE, energy, SD, CSD, GSD and Gaussian MMD, with sliced *W*_1_ also evaluated on the circle and sinusoidal tasks. MSE uses the sampled batch pairing. Architectures and budgets are fixed within each task; hyperparameters are not tuned against held-out outcomes.

**Fig. S3:**
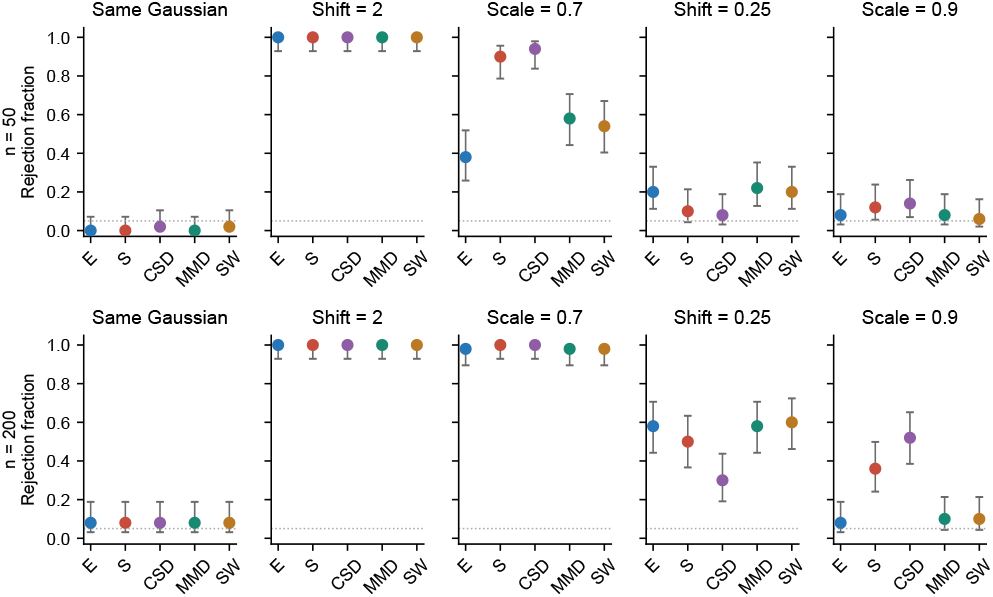
Complete sensitivity comparison: 50 independent sample pairs for each sample size and scenario, 999 pooled permutations per pair. Points are per-test rejection fractions at 0.05; bars are 95% Wilson intervals. E and S are raw statistics.

**Fig. S4:**
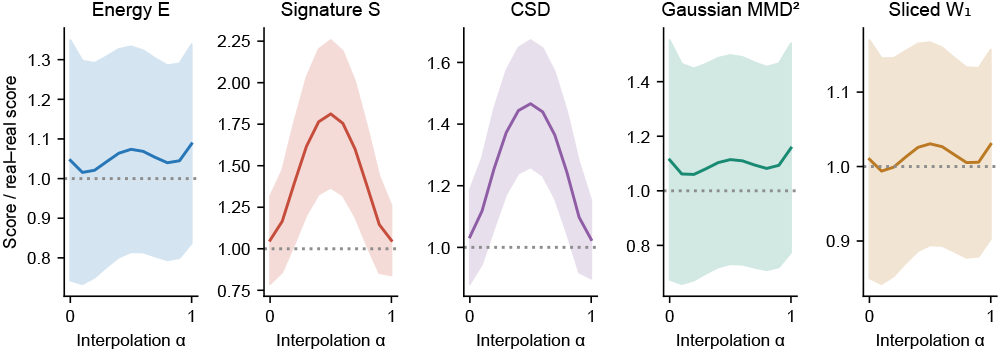
Interpolation against an independent reference population over 20 patient-disjoint partitions. Each statistic is divided by its real–real reference value; energy and signature ratios use raw ε and S. Bands show one standard deviation across partitions. Main Figure 4 compares the interpolants directly with their destination population.

The synthetic experiments optimize ε, S, CSD and GSD as defined in the main text. All networks use Adam; generator minibatches contain 256 observations. Shared architecture and optimizer settings are summarized below.

| Network | Hidden layers | Learning rate | Budget |
| --- | --- | --- | --- |
| Circle | $5 \times 128$ | $10^{-3}$ | 10,000 updates |
| Sinusoid | $2 \times 128$ | $3 \times 10^{-4}$ | 7,000 updates |
| TCGA generator | $5 \times 512$ | $10^{-3}$ | 100,000 updates |
| Classifier | $3 \times 256$ | $10^{-3}$ | 500 epochs |

Gaussian MMD uses a fixed median-distance bandwidth estimated from the training data. The sliced *W*_1_ loss is the mean absolute difference between sorted projections of generated and real observations, averaged over 128 directions. Unit directions are obtained by normalizing isotropic Gaussian vectors and are shared across updates and training runs; evaluation uses an independent direction bank.

#### Synthetic generation

All seven loss functions use three independent training runs per synthetic task. Tables report means ± standard deviations; the sample plots show a prespecified run.

**Fig. S5:**
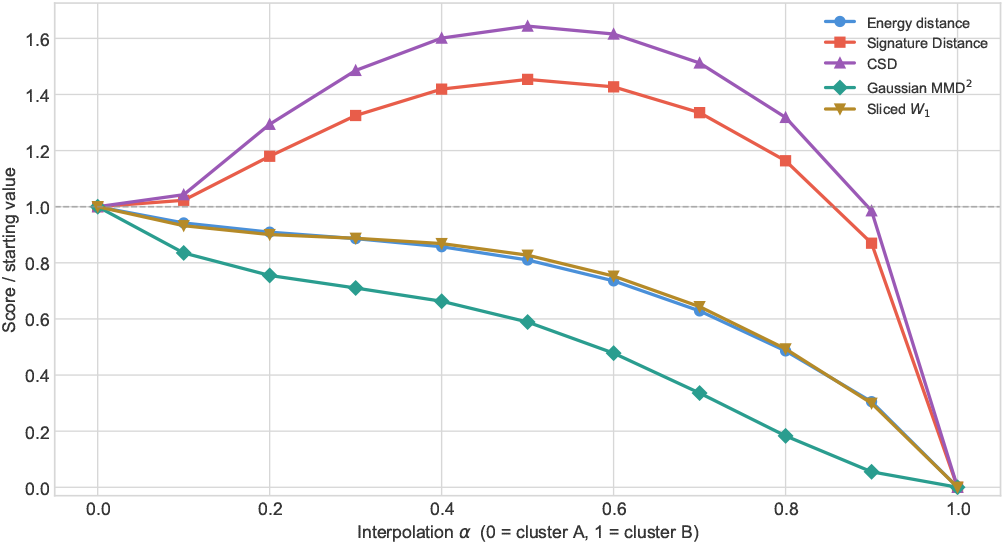
Five-method comparison on the interpolation experiment in main Figure 4. The 400-specimen populations A and B and their optimal matching are shared across methods. Scores are divided by their starting values; the grey dashed line marks one. ED and SD are rooted; CSD, Gaussian MMD^2^ and sliced W_1_ retain their stated conventions. MMD uses the median positive distance of pooled A and B as its fixed bandwidth. Sliced W_1_ uses 128 fixed projection directions. All five scores are zero at *α*=1, where the compared empirical distributions coincide.

For the two-circle task, the target has 2,000 observations per class: radius 2 centred at (−1, 0) and radius 3.5 centred at (2.5, 0), with independent coordinate noise of standard deviation 0.08. Inputs are standardized using the fixed training sample. The multilayer perceptron (MLP) generator uses rectified linear unit (ReLU) activations, two latent Gaussian coordinates and the class indicator concatenated at each layer. Updates alternate between classes; all runs share the same target sample.

| Loss | Sliced $W_1$ | Precision | Recall | Radial error |
| --- | --- | --- | --- | --- |
| MSE | $0.584 \pm 0.003$ | $0.495 \pm 0.001$ | $0.001 \pm 0.001$ | $2.679 \pm 0.021$ |
| ED | $0.048 \pm 0.016$ | $0.783 \pm 0.074$ | $0.936 \pm 0.046$ | $0.144 \pm 0.034$ |
| SD | $0.044 \pm 0.015$ | $0.861 \pm 0.031$ | $0.916 \pm 0.056$ | $0.116 \pm 0.011$ |
| CSD | $0.048 \pm 0.014$ | $0.869 \pm 0.051$ | $0.915 \pm 0.038$ | $0.098 \pm 0.021$ |
| GSD | $0.035 \pm 0.008$ | $0.970 \pm 0.018$ | $0.619 \pm 0.039$ | $0.050 \pm 0.011$ |
| MMD | $0.084 \pm 0.012$ | $0.321 \pm 0.021$ | $0.979 \pm 0.006$ | $0.608 \pm 0.034$ |
| Sliced $W_1$ | $0.047 \pm 0.015$ | $0.852 \pm 0.049$ | $0.889 \pm 0.062$ | $0.114 \pm 0.018$ |

For the heteroscedastic sinusoid, *t* ∼ *U* (0, 2*π*) and *y* = sin *t* + *σ*(*t*)*ξ*, with *σ* = 0.06 below *π* and 0.28 above, and independent standard Gaussian *ξ*. The generator takes (*t, ϵ*) and learns both output coordinates (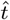, *ŷ*), using Gaussian error linear unit activations. Training uses 2,000 observations. Evaluation shares 4,000 held-out input *t* values between the real observations and generator inputs, with fresh latent noise. The real–real control uses an independently drawn second set of covariates.

| Loss | Sliced $W_1$ | Precision | Recall |
| --- | --- | --- | --- |
| MSE | $0.028 \pm 0.004$ | $0.986 \pm 0.003$ | $0.213 \pm 0.072$ |
| ED | $0.018 \pm 0.001$ | $0.969 \pm 0.005$ | $0.862 \pm 0.013$ |
| SD | $0.018 \pm 0.002$ | $0.983 \pm 0.001$ | $0.699 \pm 0.036$ |
| CSD | $0.019 \pm 0.003$ | $0.975 \pm 0.003$ | $0.764 \pm 0.046$ |
| GSD | $0.020 \pm 0.001$ | $0.983 \pm 0.001$ | $0.641 \pm 0.052$ |
| MMD | $0.042 \pm 0.004$ | $0.908 \pm 0.027$ | $0.893 \pm 0.005$ |
| Sliced $W_1$ | $0.017 \pm 0.002$ | $0.978 \pm 0.001$ | $0.782 \pm 0.026$ |

#### Sine residual spread

| Loss | Narrow SD | Wide SD |
| --- | --- | --- |
| MSE | $0.013 \pm 0.003$ | $0.018 \pm 0.002$ |
| ED | $0.052 \pm 0.000$ | $0.273 \pm 0.003$ |
| SD | $0.029 \pm 0.002$ | $0.225 \pm 0.007$ |
| CSD | $0.042 \pm 0.005$ | $0.249 \pm 0.009$ |
| GSD | $0.022 \pm 0.004$ | $0.194 \pm 0.007$ |
| MMD | $0.103 \pm 0.011$ | $0.367 \pm 0.023$ |
| Sliced $W_1$ | $0.038 \pm 0.003$ | $0.255 \pm 0.005$ |

The sine residual is generated *y* minus sin 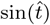, partitioned at generated 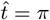. The nominal target residual standard deviations are 0.06 and 0.28. These joint-distribution diagnostics distinguish geometric proximity from recovery of the target noise scales.

#### TCGA generation

The TCGA generator uses 32 latent Gaussian coordinates and a 24-dimensional one-hot label, with layer normalization and leaky ReLU; the input is concatenated at every hidden layer. The global/local objective averages global and within-label losses (Equation 10), including labels represented by at least two samples in the batch. ED and SD optimize 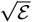; and 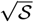; CSD, GSD and MSE use their stated definitions.

Validation uses four fixed batches of 256 observations with fixed latent noise, shared across objectives within each run. It is checked every 25 updates, with patience of 200 checks. The lowest-validation-loss state is retained. The global-only control uses MSE, ε, *S*, CSD and GSD on the same patient partitions, architecture and maximum budget, omitting within-label losses. It uses fresh validation observations and noise every five updates, with patience of 1,000 checks. The Gaussian MMD global/local comparison uses this same stochastic-validation schedule.

Each TCGA objective and batch protocol uses five independent training runs on fixed patient partitions. The classifier uses ReLU and layer normalization, and is trained on synthetic observations numbering ten times the combined fitting/validation pool, with matching label proportions. Early stopping uses a 20% synthetic validation split and patience of 50 epochs. Classification accuracy and macro-F1 are evaluated on real test observations from held-out patients.

Independent geometric precision and recall use the *k* = 3 nearest-neighbour-ball construction (Kynkäänniemi et al., 2019; main reference list), calculated in all 978 standardized features. Generated and real test populations have equal sample budgets, with a real–real comparison for context. Sliced *W*_1_ uses 128 fixed directions; label-specific diagnostics use up to 256 points per side. Reported standard deviations quantify training and sampling variation at the fixed patient split.

The global/local geometric results complement main Table 1. Values are means ± standard deviations across five independent training runs on the fixed patient split.

**Table 1.** Tissue-conditioned generation on TCGA landmark genes (978 genes, 24 tissues, noise dim = 32). All losses are trained with the same glocal protocol (Equation 10), ensuring fair comparison; only the loss *D* varies. Values are means *±* standard deviations over five seeds; bold indicates the best mean per column. MMD and independent generation diagnostics are in the Supporting Information.

| Loss | Acc (%) | F1 (%) |
| --- | --- | --- |
| MSE | 85.59 ± 0.87 | 79.93 ± 0.93 |
| ED | 92.21 ± 0.33 | **88.63 ± 0.47** |
| SD | 90.71 ± 0.63 | 87.25 ± 0.68 |
| CSD | 91.34 ± 0.92 | 87.72 ± 1.10 |
| GSD | **92.66 ± 0.43** | 88.59 ± 0.35 |

| Loss | Precision | Recall | Sliced $W_1$ |
| --- | --- | --- | --- |
| MSE | $1.000 \pm 0.000$ | $0.000 \pm 0.000$ | $0.395 \pm 0.006$ |
| ED | $0.369 \pm 0.049$ | $0.126 \pm 0.018$ | $0.076 \pm 0.005$ |
| SD | $0.126 \pm 0.014$ | $0.192 \pm 0.024$ | $0.083 \pm 0.006$ |
| CSD | $0.134 \pm 0.009$ | $0.202 \pm 0.033$ | $0.081 \pm 0.005$ |
| GSD | $0.498 \pm 0.065$ | $0.119 \pm 0.020$ | $0.082 \pm 0.005$ |

The Gaussian MMD comparison uses stochastic validation: accuracy 0.884 ± 0.014, macro-F1 0.816 ± 0.021, precision 0.644 ± 0.149, recall 0.144 ± 0.156 and sliced W_1_ 0.177 ± 0.053.

**Fig. S6:**
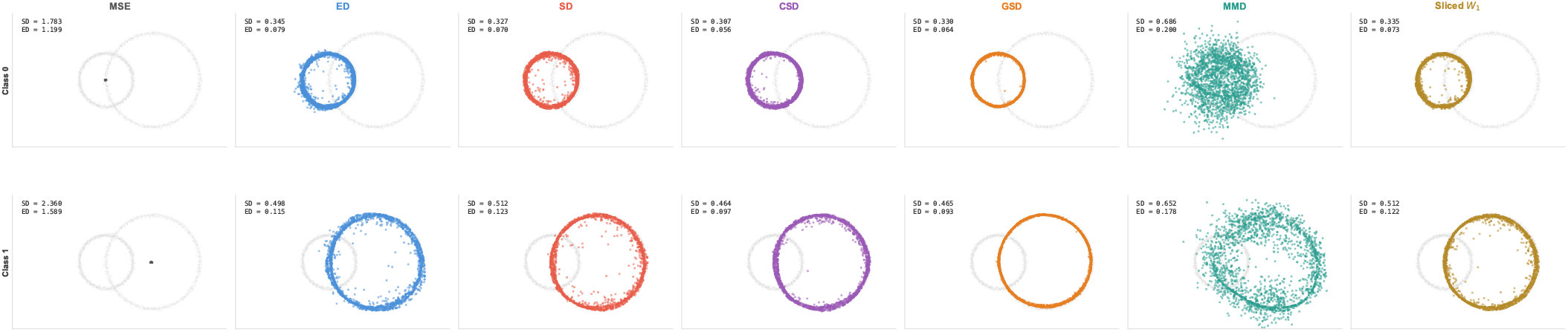
Circle generation with seven loss functions, including Gaussian MMD and sliced W_1_. Each column is a loss and each row a conditioning class; grey points show the training distribution. Labels report SD and ED against independent real samples in the original coordinate system. Repeated evaluations are reported in Section S3.

**Fig. S7:**
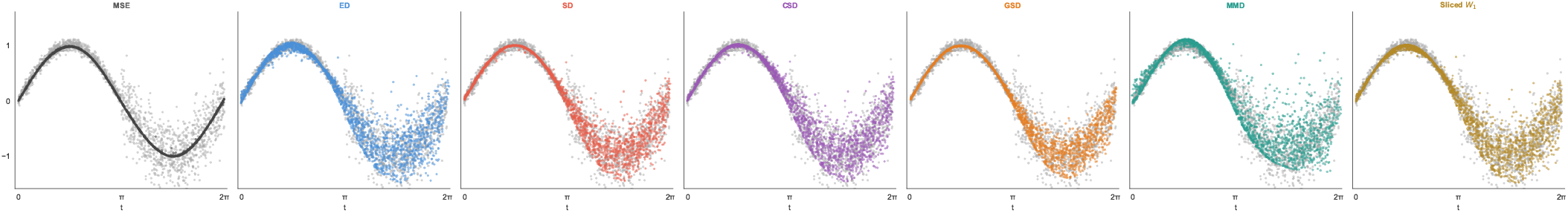
Sinusoidal generation with seven loss functions, including Gaussian MMD and sliced W_1_. The display shows 2,000 generated and held-out target observations from a prespecified run. Repeated residual-spread results use the full 4,000-point evaluation sample (Section S3).

#### Global-only control

| Loss | Accuracy | Macro F1 | Precision | Recall | Sliced $W_1$ |
| --- | --- | --- | --- | --- | --- |
| MSE | $0.850 \pm 0.007$ | $0.795 \pm 0.006$ | $1.000 \pm 0.000$ | $0.000 \pm 0.000$ | $0.400 \pm 0.004$ |
| ED | $0.866 \pm 0.006$ | $0.823 \pm 0.008$ | $0.343 \pm 0.041$ | $0.055 \pm 0.010$ | $0.068 \pm 0.006$ |
| SD | $0.565 \pm 0.236$ | $0.403 \pm 0.178$ | $0.046 \pm 0.007$ | $0.166 \pm 0.007$ | $0.087 \pm 0.009$ |
| CSD | $0.366 \pm 0.212$ | $0.256 \pm 0.160$ | $0.059 \pm 0.013$ | $0.209 \pm 0.018$ | $0.087 \pm 0.007$ |
| GSD | $0.164 \pm 0.022$ | $0.127 \pm 0.018$ | $0.277 \pm 0.027$ | $0.071 \pm 0.014$ | $0.099 \pm 0.011$ |

The global-only comparison uses the same patient partitions and seeds, with raw energy and signature objectives and stochastic validation.

#### Population expansion

The general RNA-seq representation selects the 256 highest-variance features. A 10-nearest-neighbour graph and Leiden resolution 1 produce 21 clusters; the largest contains 931 patients. A partition reserves 310 observations each for fitting A, validation B and test C, leaving one observation unused. Feature selection and clustering precede this partition. Five bootstrap resamples of A are each initialized from 31 k-means centres, replicated by cluster occupancy to produce 310 candidate points. The update uses step size 0.1 and noise amplitude 0.1 for 500 steps; coordinate values are clipped below at zero. Noise follows the regularized bootstrap covariance, with its Cholesky factor normalized by 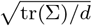, and is shared across objectives within each bootstrap. The gradient is computed with respect to the candidate points, in chunks of at most 256, against the complete fixed bootstrap reference. Each chunk is treated as the generated empirical sample for its guidance objective; within-generated distances are computed within that chunk. Stopping and reported distances use the complete candidate population. Guidance uses ε, S or CSD. The stopping score in the main text uses rooted energy distance for energy guidance and rooted SD for SD/CSD guidance. It is evaluated on the complete candidate population before each update. Among the five trajectories for each objective, the main figure displays the one with lowest validation distance at its criterion-selected stopping step. Test C is used only for the reported traces and independent evaluation, never for stopping or replicate selection. The PCA is fitted on the full selected 931-patient cluster. Initial-centroid markers correspond to the displayed bootstrap trajectory.

| Loss | Step | Precision | Recall | Sliced $W_1$ |
| --- | --- | --- | --- | --- |
| ED | $174.2 \pm 71.6$ | $0.802 \pm 0.134$ | $0.208 \pm 0.180$ | $0.321 \pm 0.028$ |
| SD | $217.8 \pm 11.6$ | $0.721 \pm 0.021$ | $0.293 \pm 0.046$ | $0.317 \pm 0.027$ |
| CSD | $220.0 \pm 10.3$ | $0.716 \pm 0.027$ | $0.299 \pm 0.045$ | $0.318 \pm 0.027$ |

Each row summarizes five bootstrap trajectories. Independent precision, recall and sliced *W*_1_ are evaluated on the reserved test sample after stopping.

#### Normalization and component illustrations

The exploratory normalization and component comparison in Figure 9 uses leave-one-out within-reference profiles and interpolated cross-profile ranks. NSD rescales coordinates by the detached median cross-distance before computing the raw signature loss. This differs from SD_norm_ in Methods, whose denominator uses the mean cross-distance. The repeated experiments in Section S3 use the complete empirical estimator in Appendix C.

## Footnotes

1 https://www.cancer.gov/tcga

## Notes

https://github.com/lazzaronico/signature-distance

### Summary of Updates

We adopt the summed convention S = SDiv(X‖Y) + SDiv(Y‖X), with SD = √S, so that E ≤ S ≤ 2W₁ and S reduces to the standard raw energy statistic E under the stated radial stochastic-dominance condition. This normalization rescales raw values by two and rooted values by √2. Separately, the statistical implementation and repeated experiments now retain complete empirical atoms and integrate unequal-size radial profiles exactly. The main global/local TCGA comparison optimizes rooted ED and SD; synthetic, global-only TCGA and expansion comparisons optimize raw E and S. CSD/GSD retain their definitions without compensating multipliers.

## References

Argelaguet R, Velten B, Arnol D, et al. Multi-Omics Factor Analysis — a framework for unsupervised integration of multiomics data sets. Molecular Systems Biology. 2018;14(6):e8124.

Beyer K, Goldstein J, Ramakrishnan R, Shaft U. When is “nearest neighbor” meaningful? Database Theory — ICDT’99, Lecture Notes in Computer Science. 1999;1540:217–235.

Carlsson G. Topology and data. Bulletin of the American Mathematical Society. 2009;46(2):255–308.

Chawla NV, Bowyer KW, Hall LO, Kegelmeyer WP. SMOTE: synthetic minority over-sampling technique. Journal of Artificial Intelligence Research. 2002;16:321–357.

Cui H, Wang C, Maan H, et al. scGPT: toward building a foundation model for single-cell multi-omics using generative AI. Nature Methods. 2024;21(8):1470–1480.

Cuturi M. Sinkhorn distances: lightspeed computation of optimal transport. Advances in Neural Information Processing Systems. 2013;26:2292–2300.

Dixit A, Parnas O, Li B, et al. Perturb-Seq: dissecting molecular circuits with scalable single-cell RNA profiling of pooled genetic screens. Cell. 2016;167(7):1853–1866.

Edelsbrunner H, Harer JL. Computational Topology: An Introduction. American Mathematical Society; 2010.

Efron B, Tibshirani RJ. An Introduction to the Bootstrap. Chapman & Hall/CRC; 1993.

Ernst MD. Permutation methods: a basis for exact inference. Statistical Science. 2004;19(4):676–685.

Goodfellow I, Pouget-Abadie J, Mirza M, et al. Generative adversarial nets. Advances in Neural Information Processing Systems. 2014;27:2672–2680.

Gretton A, Borgwardt KM, Rasch MJ, Schölkopf B, Smola AJ. A kernel two-sample test. Journal of Machine Learning Research. 2012;13:723–773.

Heusel M, Ramsauer H, Unterthiner T, Nessler B, Hochreiter S. GANs trained by a two time-scale update rule converge to a local Nash equilibrium. Advances in Neural Information Processing Systems. 2017;30.

Kingma DP, Welling M. Auto-encoding variational Bayes. International Conference on Learning Representations. 2014.

Kuhn HW. The Hungarian method for the assignment problem.Naval Research Logistics Quarterly. 1955;2(1–2):83–97.

Kynkäänniemi T, Karras T, Laine S, Lehtinen J, Aila T. Improved precision and recall metric for assessing generative models. Advances in Neural Information Processing Systems. 2019;32.

Lloyd S. Least squares quantization in PCM. IEEE Transactions on Information Theory. 1982;28(2):129–137.

Lopez R, Regier J, Cole MB, Jordan MI, Yosef N. Deep generative modeling for single-cell transcriptomics. Nature Methods. 2018;15(12):1053–1058.

Luecken MD, Theis FJ. Current best practices in single-cell RNA-seq analysis: a tutorial. Molecular Systems Biology. 2019;15(6):e8746.

Marouf M, Machart P, Bansal V, et al. Realistic in silico generation and augmentation of single-cell RNA-seq data using generative adversarial networks. Nature Communications. 2020;11(1):166.

Peyré G, Cuturi M. Computational optimal transport.Foundations and Trends in Machine Learning. 2019;11(5–6):355–607.

Phipson B, Smyth GK. Permutation P-values should never be zero: calculating exact P-values when permutations are randomly drawn. Statistical Applications in Genetics and Molecular Biology. 2010;9(1):Article 39.

Rezende DJ, Mohamed S. Variational inference with normalizing flows. International Conference on Machine Learning. 2015:1530–1538.

Subramanian A, Narayan R, Corsello SM, et al. A next generation connectivity map: L1000 platform and the first 1,000,000 profiles. Cell. 2017;171(6):1437–1452.

Székely GJ, Rizzo ML. Energy statistics: A class of statistics based on distances. Journal of Statistical Planning and Inference. 2013;143(8):1249–1272.

The Cancer Genome Atlas Research Network. Comprehensive genomic characterization defines human glioblastoma genes and core pathways. Nature. 2008;455(7216):1061–1068.

Theodoris CV, Xiao L, Chopra A, et al. Transfer learning enables predictions in network biology. Nature. 2023;618(7965):616–624.

Traag VA, Waltman L, van Eck NJ. From Louvain to Leiden: guaranteeing well-connected communities. Scientific Reports.2019;9:5233.

Villani C. Optimal Transport: Old and New. Springer; 2009.

Welling M, Teh YW. Bayesian learning via stochastic gradient Langevin dynamics. International Conference on Machine Learning. 2011:681–688.

Sejdinovic D, Sriperumbudur B, Gretton A, Fukumizu K. Equivalence of distance-based and RKHS-based statistics in hypothesis testing. Annals of Statistics. 2013;41(5):2263–2291.

Rodriguez A, Laio A. Clustering by fast search and find of density peaks. Science. 2014;344(6191):1492–1496.

